# Sleep, NMDA Receptor Subunits, and the Compensatory Pathway: Understanding Contextual Fear Conditioning in the Absence of the Dorsal Hippocampus

**DOI:** 10.1101/2024.06.07.597897

**Authors:** Deepika Kant, Sushil K. Jha

## Abstract

The loss of the dorsal hippocampus (DH) results in profound deficits in contextual fear-conditioned (CxFC) memory. Nonetheless, CxFC memories can still form without the DH, specifically with multiple trials at three-day intervals. The infralimbic cortex (IL) is pivotal in initiating this compensatory process post-DH loss, but the precise factors remain elusive. Our study aims to delineate key factors of compensatory CxFC in DH absence by investigating the effects of sleep deprivation (SD) and NMDA receptor subunits NR2A and NR2B. Using a DH-lesioned rat model, we conducted two conditioning trials separated by three days and assessed fear response during the subsequent test. We observed that DH-lesioned animals exhibited to SD (DHL-SD) did not elicit a compensatory CxFC response, displaying significantly impaired freezing during the second test. Conversely, DH-lesioned non-sleep-deprived animals (DHL-NSD) compensated for DH loss and exhibited robust CxFC responses during the second test. Moreover, inhibiting NR2B subunits in the IL during initial CxFC training disrupted the formation of compensatory fear memory in DH-lesioned animals, while NR2A subunit blockade showed no significant effect. These emphasize the adverse impact of SD on compensatory memory and the critical role of NR2B subunits in facilitating compensatory CxFC memory formation following DH loss.

## Introduction

In memory processes, the adaptability of neural circuits is a central question. The encoding and retention of memories rely on intricate interactions among various brain regions. The dorsal hippocampus (DH) is crucial in consolidating and retrieving memories like contextual fear memory. Loss of DH function results in profound impairments in such memory (Matus-Amat *et al*., 2004; Wiltgen *et al*., 2006; Zelikowsky *et al*., 2013). However, studies suggest alternative neural circuits can compensate for the DH loss. Additional training trials can gradually develop contextual fear within these alternative circuits, indicating their remarkable adaptive potential (Coelho *et al*., 2018; Cowansage *et al*., 2014; Kant and Jha, 2019a, 2023; M. S. Fanselow, 2010; Poulos *et al*., 2010; Yu *et al*., 2021; Zelikowsky *et al*., 2013). In the absence of the DH, the micro-circuitries of the medial prefrontal cortex (mPFC), particularly the infralimbic (IL) and prelimbic (PL) cortices, facilitate contextual fear conditioning (Zelikowsky *et al*., 2013). Furthermore, recent findings indicate that compensatory contextual fear memory can develop following multiple conditioning trials separated by three days, even in the absence of the DH (Kant and Jha, 2019b, 2023). Notably, the input of contextual fear memory to the IL during the initial conditioning session is crucial for inducing and progressing the compensatory network in the absence of DH (Kant and Jha, 2023). However, beyond timing and extensive training, other factors influencing compensatory contextual fear memory processes in alternative brain circuitry remain to be explored.

Sleep plays a significant role in developing, maturing, and remodeling neural circuits, crucial for synaptic plasticity and functional connectivity across species and brain regions (Cao *et al*., 2020; Sun *et al*., 2020; Weiss and Donlea, 2021). Research highlights sleep’s involvement in rapid synapse addition in young drosophila’s olfactory ganglia development and ocular dominance plasticity in cats, where sleep promotes twice as many new brain connections after environmental stimulation (Jha *et al*., 2005a; Kayser *et al*., 2014). Moreover, sleep regulates critical periods and synaptic plasticity in the visual cortex, strengthening and maintaining newly formed spines, thus enhancing learning performance (Frank, 2015; Frank *et al*., 2001). Conversely, a lack of sleep disrupts long-term potentiation (LTP) stability and glutamatergic neurotransmitter signaling, impairing hippocampal neuronal circuit maturation (Grønli *et al*., 2014; Lopez *et al*., 2008). Sleep deprivation negatively affects various memory processes, including spatial memory, delay-conditioned memory, cued fear memory, and contextual fear memory (Graves *et al*., 2003; Qureshi and Jha, 2017; T. Kumar and S. K. Jha, 2017; Tripathi and Jha, 2016; Ward *et al*., 2017).

N-methyl-D-aspartate receptors (NMDARs) are pivotal for LTP and memory plasticity in the brain, particularly in fear learning and contextual memory formation (Baez *et al*., 2018; Wang and Peng, 2016). Comprising mandatory NR1 and optional NR2 subunits, NMDARs have isoforms NR2A-D with distinct physiological properties. NR2A and NR2B subunits, highly expressed in fear-regulating brain regions like the hippocampus, prefrontal cortex, and amygdala, play region-specific roles in contextual fear memory expression (Liu *et al*., 2004). While NR2B in the anterior cingulate cortex is crucial for both LTP and contextual fear memory, NR2B in the PL mediates trace conditioning alone (Zhao *et al*., 2005). Conversely, NR2A in PL regulates both trace and contextual fear memory (Gilmartin *et al*., 2013). However, the precise contribution of NR2A and NR2B subunits in the formation of compensatory fear memory mediated by the IL remains unclear. Our study aims to elucidate the impact of chronic sleep deprivation and selective NR2A/NR2B blockade in the IL on compensatory contextual fear memory formation in the absence of the DH.

## Methodology

All the experiments were carried out on male Wistar rats (230-270 g, 10-12 weeks old), brought from the university’s “Central Laboratory Animal Resources” (CLAR). Animals were maintained on a 12:12 light-dark cycle (lights on from 7:00 AM to 7:00 PM) in the air-conditioned environment at 20–22 °C ambient room temperatures. Food and water were given *ad libitum.* We have also taken utmost care to minimize the use of animals in experiments. All procedures were approved by the Institution’s Animal Ethical Committee (IAEC protocol # 22/2014) of Jawaharlal Nehru University, New Delhi.

Experiment I: we investigated the significance of sleep in the development of compensatory contextual fear memory after DH loss. Experiment II: we examined the role of NMDARs subunits NR2A and NR2B in the IL-dependent development of compensatory contextual fear memory following DH loss.

Animals underwent stereotaxic surgery for drug microinjection using a previously established protocol. Briefly, stainless steel cannulae (24-gauge; 15 mm length) were implanted bilaterally in the DH region (−2.8 mm AP, ±1.6 mm ML, and 2.5 mm DV; −4.2 mm AP, −2.6 mm ML, and 2.5 mm DV) for ibotenic acid microinjection at four sites, causing maximum neuronal damage. For Experiment II, two additional cannulae were implanted bilaterally in the IL region (+3.0 mm AP, ±0.6 mm ML, and 4.0 mm DV). Following surgery, the animals were provided with post-op care and allowed one week’s recovery before experiments.

### Experimental design

#### Experiment I - Sleep and Compensatory Contextual Fear Memory

We investigated the effect of sleep on the induction of compensatory pathways for contextual fear memory in the absence of the DH. We designed the experiments based on our previously established protocol (Kant and Jha, 2023) where a compensatory contextual fear memory is formed in DH-lesioned animals through two contextual fear conditioning (CxFC) sessions separated by a 3-day interval (72-h). On day 1, baseline freezing was recorded in surgically prepared animals, after 2 d of habituation in the neutral behavioral chamber. Thereafter, the DH-lesioned group received microinjections of ibotenic acid (IBA; an excitotoxin), while the DH-non-lesioned group received vehicle (PBS) microinjections. After 42 h, both groups underwent the first CxFC training on day 3 and tested on day 4. Animals {DH-lesioned sleep-deprived (DHL-SD; n=8) and DH-non-lesioned sleep-deprived (DHNL-SD; n=8)}group were then subjected to 54 h of sleep deprivation, followed by 18 h of recovery sleep, creating a 3-day (72-hour) period. Additionally, a DH-non-lesioned non-sleep-deprived (DHNL-NSD; n=7) group was kept in their home cages during the equivalent 72-hour period. Animals received 2^nd^ CxFC training on day 8 and tested (testing 2) on day 9 (Fig. 2).

#### Experiment II - NMDAR Subunits NR2A and NR2B in Compensatory Contextual Fear Memory

We investigated the role of NMDAR subunits NR2A and NR2B in the infralimbic cortex (IL)-mediated compensatory contextual fear memory in DH-lesioned animals. After IBA microinjection, DH-lesioned animals were divided into three groups: (1) NR2A-containing NMDA receptor antagonist (PEAQX) microinjection (n=8), (2) NR2B-containing NMDA receptor antagonist (Ro25-6981) microinjection (n=8) and (3) Control group receiving vehicle (0.9% saline) microinjection (n=7). After 30 mins, animals underwent 1^st^ CxFC training on day 3 (training 1) and tested on day 4 (testing 1). Thereafter, animals underwent a 2^nd^ CxFC training (training 2) on day 8 and tested (testing 2) on day 9 (Fig. 2).

### Partial sleep deprivation using the Flowerpot method

The small flower pot platform, with a diameter of 6.5 cm, was used for sleep deprivation (Hicks *et al*., 1977). The flower pots used were plastic beakers with a capacity of 250 ml, measuring 10 cm in height and 6.5 cm in diameter. The beaker was filled with soil, and a sturdy plastic bag was used to cover the top. A plastic rope was tightly knotted around its neck to prevent water from leaking into the beaker filled with soil. In the center of a bucket partially filled with water, the beaker was kept upside down. The height of the bucket was about 35 cm and water was filled in it up to 2 cm. For sleep deprivation, the animal was placed on top of the platform kept inside the bucket, and covered with a grid. Food pellets and water bottles were provided on the holding pockets of the grid and were within the reach of animals in the standing position. The water in the bucket was changed every day at the same time. During the cleaning process, the animal was kept inside a dry cage and kept awake by gentle handling. Before placing the animal back on the platform, its weight was taken and health conditions were monitored. The control group of animals was housed in normal cages and kept in the same room as the sleep-deprived animals.

### Contextual Fear Conditioning (CxFC) Protocol

During the first two days (Days 1 and 2), the animal was habituated in the neutral chamber for 5 minutes each day between 10:55 and 11:00 AM. The baseline spontaneous freezing behavior was recorded on Day 3 in the neutral chamber for 5 minutes using FreezeFrame software (Coulbourn Inc, USA) and a CCTV camera (SenTech, USA). Illumination of 20 Lux was kept in the neutral chamber during the habituation and baseline recording (Days 1-3). On Day 4, the animal was brought to the behavioral chamber by an unfamiliar person via a different route from the animal room. To distinguish between the neutral and shock chambers, situational cues, such as an illumination of 80 Lux and a sandalwood fragrance soaked in a cotton ball, were incorporated. The same husk bedding used during fear-conditioning training was used again during testing. During training, three scrambled foot shocks (0.8 mA; 2 s) were administered to the animal at 1-minute intervals over 5 minutes using FreezeFrame software (Coulbourn Inc, USA; model #H13-17) through the grid floor of the shock chamber. The animal’s freezing behavior was recorded by a CCTV camera (SenTech, USA) for 5 minutes, during a time-matched hour on baseline days, for offline analysis. On Day 5, the animal’s CxFC was tested in the same chamber with the same cues as the training day, but no foot shock was administered. The induced freezing was captured on camera for the complete 5 minutes. Statistical comparison was made between changes in the freezing response on baseline and post-conditioning days.

### Data analysis of freezing behavior

Freezing behavior was analyzed using FreezeFrame software. Freezing was recorded if the animal stayed motionless for at least 2 sec below a 10% motion index threshold. Percent freezing was calculated and analyzed statistically using Sigma Stat 12.0 software between and within groups on different days, using one-way ANOVA and repeated-measures ANOVA, respectively followed by the Tukey post hoc test. Also, we compared the group and day interaction using two-way ANOVA.

### Drugs

Ibotenic acid (IBA) (Sigma-Aldrich, USA) was dissolved in 0.1 M PBS (63 mM) and stored in 20 μL aliquots at −20°C. One hour before microinjection, the IBA was thawed to room temperature. A total of 0.5 μL of IBA or vehicle (0.1 M PBS) was microinjected at a rate of 0.25 μL/min/site into the DH using a microinjection pump (Kent Scientific, USA; model # CT 06790). To allow for maximum neuronal damage, the animal was left undisturbed for 42 hours before CxFC training.

The drug PEAQX (Sigma-Aldrich, USA) was prepared by dissolving it in 0.9% saline to yield a concentration of 1.36 µg/µl (3 mM) and stored in aliquots of approximately 20 µl at - 20°C. Before microinjection, PEAQX was thawed at room temperature and vortexed to ensure it was fully dissolved. A volume of 0.5 µl of either PEAQX or vehicle (0.9% saline) was injected at a rate of 0.5 µl/min/site. Thirty minutes following injection, all animals underwent the first CxFC training.

To prepare Ro25-6981 (Sigma-Aldrich, USA) for microinjection, it was dissolved in 0.9% saline to create a concentration of 4 µg/µl (10.64 mM), and stored in 20 µl aliquots at −20 °C. Before microinjection, the solution was thawed at room temperature and vortexed to dissolve any precipitate. Each animal received a microinjection of 0.5 µl of either Ro25-6981 or vehicle (0.9% saline), administered at a rate of 0.5 µl/min/site. Thirty minutes later, all animals were subjected to the first CxFC training.

Following the completion of the experiment, the animal was euthanized via intraperitoneal injection of thiopentone (80 mg/kg, IP) and transcardially perfused with a solution of 0.9% saline followed by 10% formaldehyde. The brain was then removed and prepared for cryosectioning. Using a cryostat (Leica, CM 1850), 40 μm sections were cut and processed for cresyl violet staining. A microscopic examination was carried out to assess the extent of the DH lesion and the site of injections around the IL. The experiments were conducted in a randomized manner, and the experimenter had access to digital files of the animal group data in a blindfolded pattern.

### Quantification of lesion

DH lesion quantification involved capturing images of each 40 μm cresyl violet-stained coronal section throughout the DH using an Olympus SZ61 light microscope and Samsung SDC-313B camera. Brain section images at five different anteroposterior levels were analyzed using ImageJ software with a software-generated grid overlay (1000 pixels per grid). Total grids within the borders of DH-lesioned animals were counted, subtracted from DH-non-lesioned animals, and divided by the total grids in the DH-non-lesioned sections, multiplied by 100 to determine DH damage percent. The lesion extent was then represented in a reconstruction diagram. We used ImageJ software to map the distribution of neuronal damage in the DH region. The software’s ROI manager was utilized to calculate mean density values from the histological sections of DH-lesioned animals. Three distinct areas within a section were analyzed: no lesion, partial lesion, and complete lesion. The mean densitometric values from these areas were then computed, with the mean density value of the complete lesion area being taken as 100% lesion. The percentage changes in the mean density values of the partial and no lesion areas were determined from the complete lesion area. Finally, we depicted a heat map showing the distribution of neuronal damage within the DH using these percentage values.

## Result

### Histological findings

Ibotenic acid microinjections led to significant bilateral destruction, affecting approximately 75% of the DH area (±4.12%). In contrast, DH-non-lesioned animals displayed intact neuronal cell bodies, as shown by cresyl violet staining (Fig. 1 and b). Heat maps in Fig 1c depict the distribution of neuronal damage within the DH. Notably, there was nearly 100% neuronal loss in the medial DH region (represented by orange color) across all experiments. In addition, experiments I and II demonstrated neuronal damage ranging from 17% to 77% and 22% to 69%, respectively, in regions adjacent to the midline and lateral side of the DH, denoted by gray and yellow colors in the heatmap.

**Figure 1:**
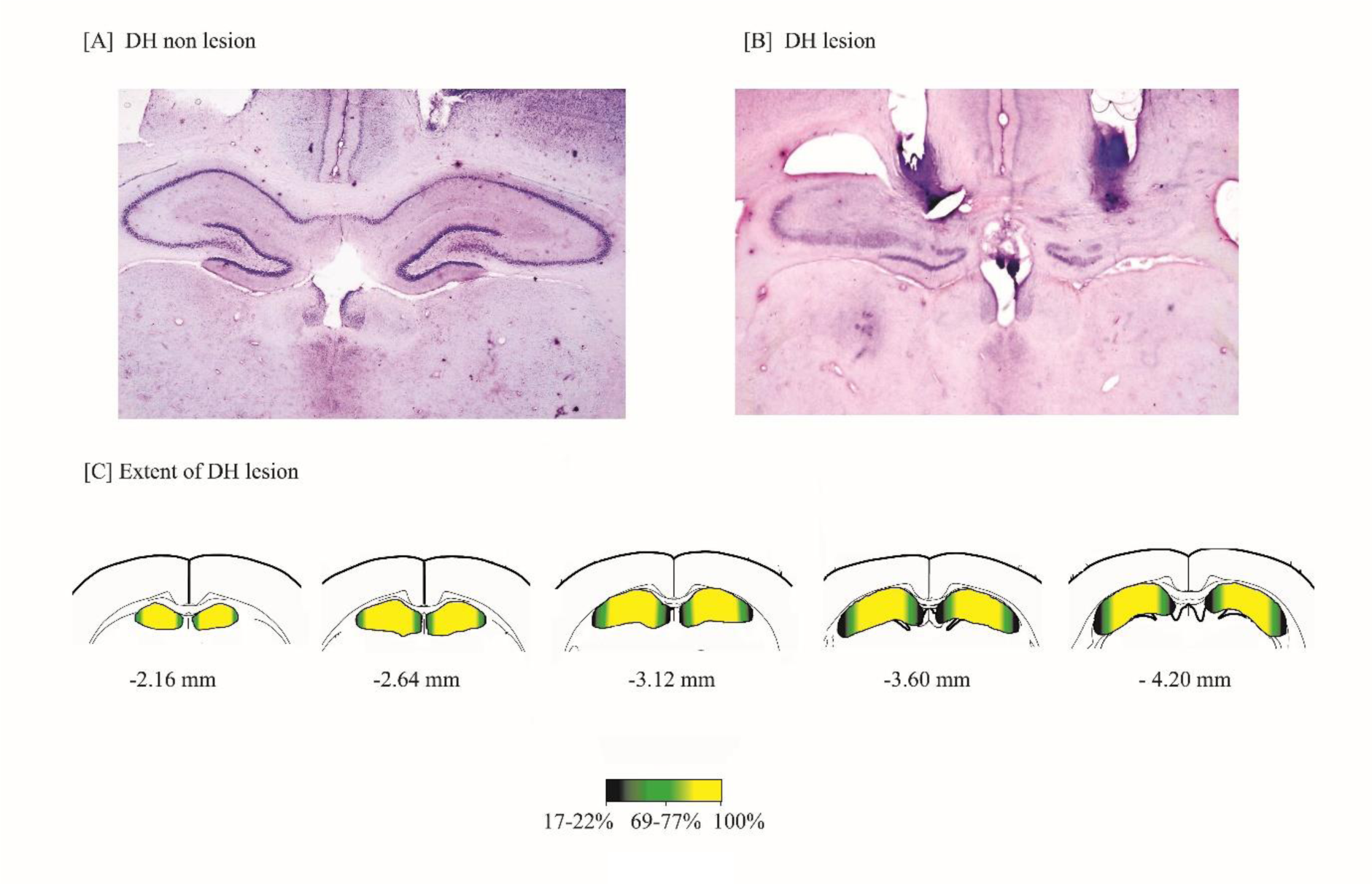
Histological microphotograph of rat brain from [A] DH-non-lesioned and [B] DH-lesioned animals, depicting intact neurons and neuronal loss, respectively. [C] The Heat map in the reconstruction diagram represents the extent of neuronal damage in DH-lesioned animals. In the heat maps, the yellow color denotes a 100% loss of neurons for both experiments I and II. Additionally, the green color denotes a 69% to 71% neuronal loss, and the black color denotes an 11% to 17% neuronal loss.

**Figure 2:**
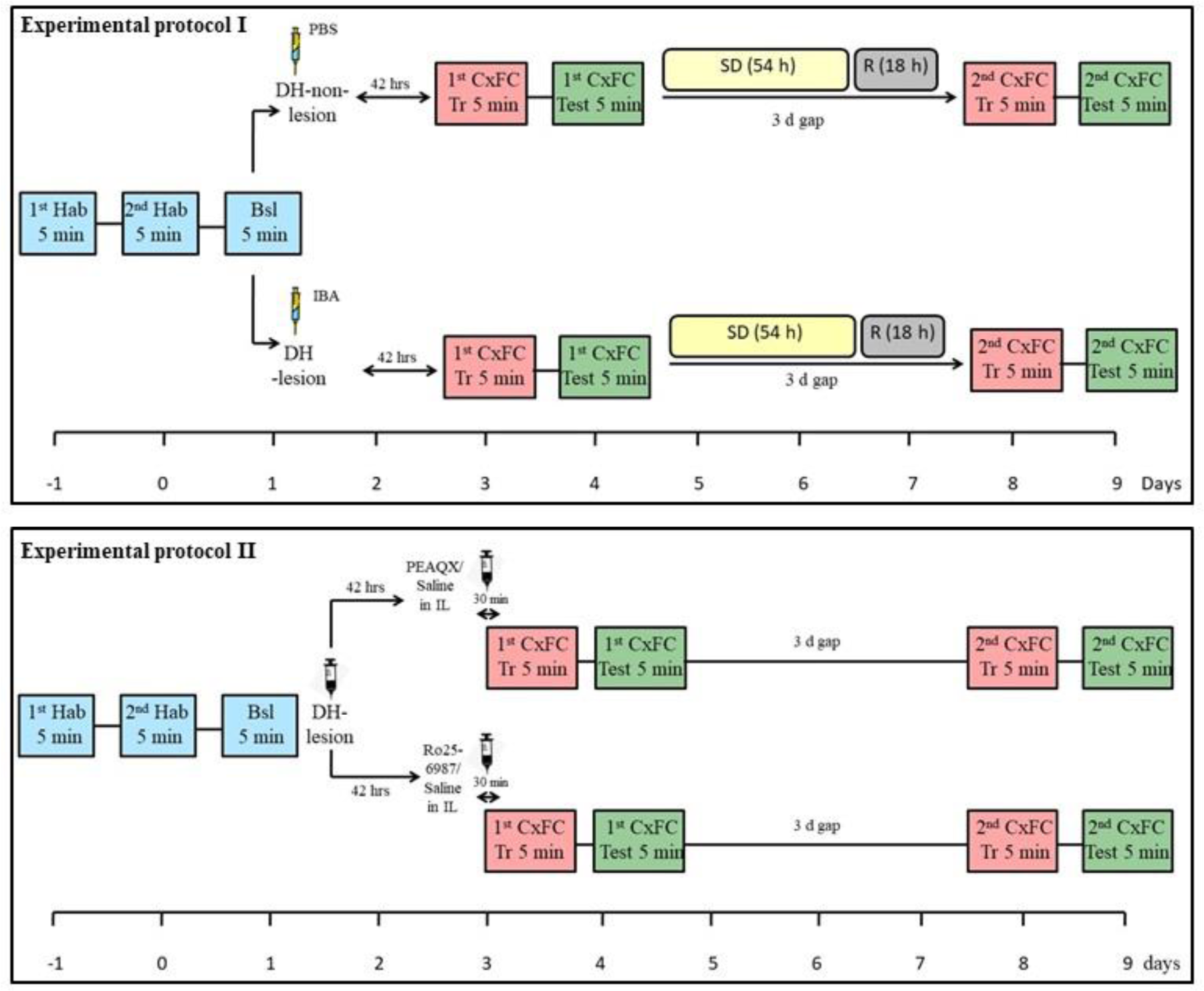
In experimental protocol I: After 2 days of habituation, baseline freezing was recorded. DH-lesioned animals received ibotenic acid (IBA) microinjections, while DH-non-lesioned animals received vehicle (PBS) microinjections. After 42 hours, both groups underwent the first contextual fear conditioning (CxFC) training on day 3, followed by testing on day 4. Subsequently, DH-lesioned and DH-non-lesioned animals were subjected to either sleep deprivation (54 hours) followed by recovery sleep (18 hours) or were kept in their home cages for an equivalent 72-hour period. The second CxFC training was conducted on day 8, followed by testing on day 9. Groups: DH-lesioned sleep-deprived (DHL-SD; n=8), DH-non-lesioned sleep-deprived (DHNL-SD; n=8), DH-non-lesioned non-sleep-deprived (DHNL-NSD; n=7). DH: Dorsal hippocampus, CxFC: Contextual fear conditioning, SD: Sleep deprivation, R: recovery sleep. In experimental protocol II: After IBA microinjection, DH-lesioned animals were divided into three groups: (1) NR2A-containing NMDA receptor antagonist (PEAQX) microinjection (n=8), (2) NR2B-containing NMDA receptor antagonist (Ro25-6981) microinjection (n=8), and (3) control group received vehicle (0.9% saline) microinjection (n=7). Animals underwent the first CxFC training on day 3, followed by testing on day 4, then a second CxFC training on day 8, followed by testing on day 9. *** denotes Tukey *p* < 0.001. Error bars represent the standard error mean (S.E.M.).

### Effect of chronic sleep deprivation (SD) on the formation of compensatory contextual fear memory in the absence of Dorsal Hippocampus (DH)

We have previously established that a three-day time interval between the two conditioning trials is essentially required for the compensatory circuitry of contextual fear to become functional after DH loss (Kant and Jha, 2019b, 2023). To address the question of whether sleep plays any role in the induction of compensatory circuitry of contextual fear, DH-lesioned animals were sleep deprived during three days or 72 h intervals (54 h of SD and 18 h of recovery sleep) between the 1^st^ and 2^nd^ CxFC trial. Details of the sleep deprivation protocols have been provided in the supplementary information.

For the changes in percent freezing in the sleep-deprived and non-sleep-deprived DH-lesioned and DH-non-lesioned animals, two-way ANOVA showed a statistically significant interaction between group and day. The post hoc comparison demonstrated that following 1^st^ CxFC trial, the DH-non-lesioned non-sleep-deprived (DHNL-NSD) animals showed significantly higher freezing response (p<0.001, F_(8,114)_ = 14.82) on the 1^st^ testing day compared to the DH-lesioned non-sleep-deprived animala (DHL-NSD) (sample variance in DHNL-NSD animals σ2 = 256.32; DHL-NSD animals σ2 = 94.61; Cohen’s d = 6.12, and power = 1.00 at the 0.05 α level) (Fig. 3). This highlights the significance of DH in contextual fear memory, as DH-lesioned animals failed to form CxFC memory.

**Figure 3:**
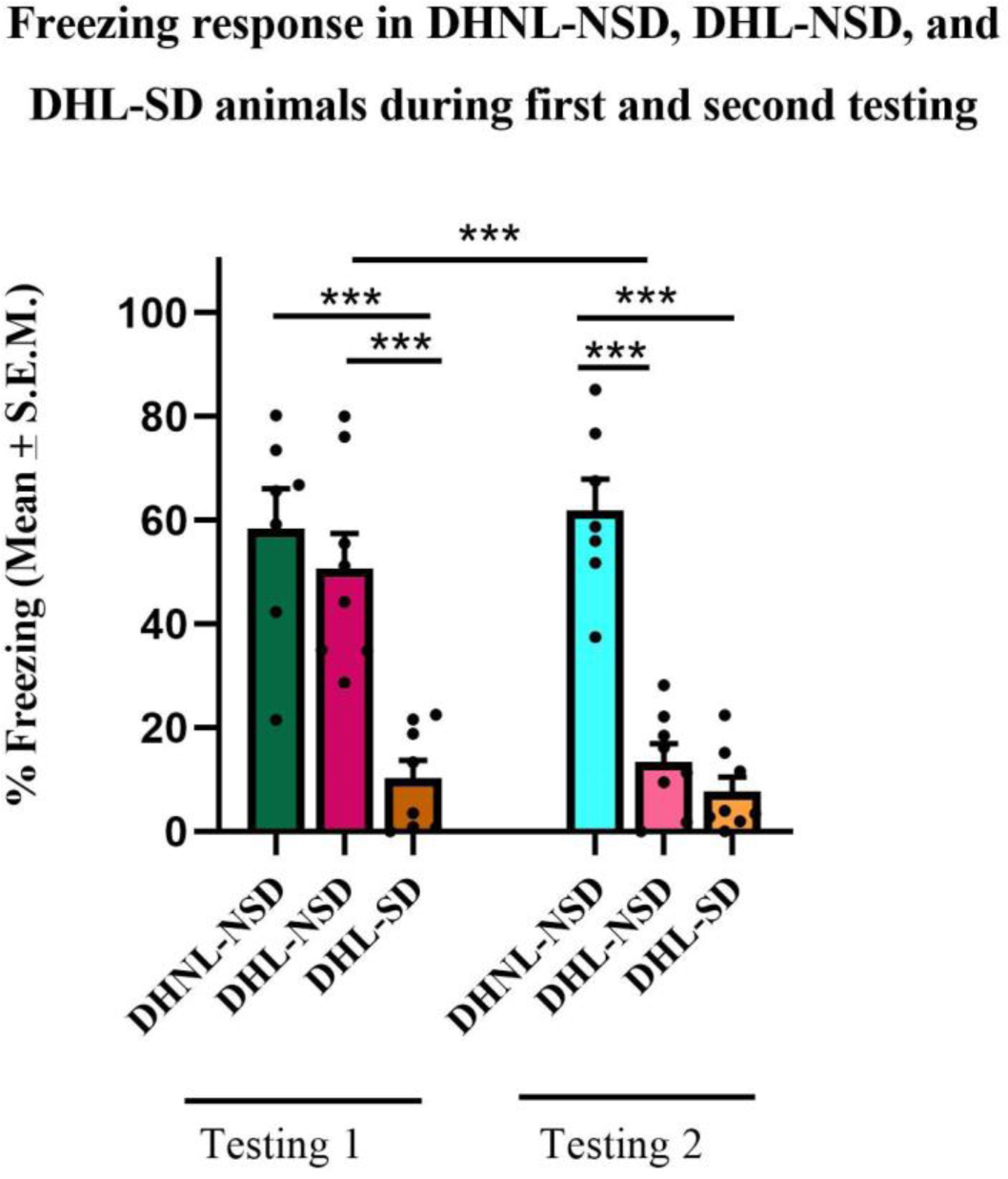
Sleep deprivation impairs compensatory contextual fear memory formation in DH-lesioned animals: DH-lesioned animals were sleep deprived during the three-day duration or 72 hours (54 h sleep deprivation and 18 h recovery sleep) between the first and second CxFC trials. A two-way ANOVA showed a significant interaction between group and day. Post hoc comparisons revealed that on the first testing day, DH-non-lesioned non-sleep-deprived (DHNL-NSD) animals exhibited a significantly higher freezing response (p<0.001) compared to DH-lesioned non-sleep-deprived (DHL-NSD) animals. After the second CxFC trial, DHL-NSD animals showed a significant increase in freezing response (p<0.001), which was comparable to DHNL-NSD animals, confirming that compensatory circuitry for contextual fear memory can develop in the absence of the DH with multiple trainings separated by three days. However, DH-lesioned sleep-deprived (DHL-SD) animals demonstrated significantly impaired freezing compared to both DHNL-NSD and DHL-NSD animals (p<0.001), highlighting that sleep deprivation impairs the formation of compensatory contextual fear memory in DH-lesioned animals. *** denotes Tukey *p* < 0.001.

After 2^nd^ conditioning trial, there was a significant increase (p<0.001) in the freezing response of the DHL-NSD group which was similar to the DHNL-NSD group (sample variance in DHL-NSD animals σ2 = 364.54; DHNL-NSD animals σ2 = 407.84; Cohen’s d = 3.72, and power = 1.00 at the 0.05 α level). This confirmed our previous finding that, in the absence of DH, functional compensatory circuitry for contextual fear memory develops through multiple trainings separated by 3 days. However, DHL-SD animals failed to compensate for the loss of DH, as indicated by their impaired freezing response on 2^nd^ testing day (Fig). Additionally, DHL-SD animals demonstrated significantly impaired freezing compared to DHNL-NSD animals (p<0.001) with an 82.59% reduction in freezing on the 2^nd^ testing day (sample variance in DHL-SD animals σ2 = 98.37; DHNL-NSD animals σ2 = 407.84; Cohen’s d = 0.39, and power = 1.00 at the 0.05 α level). Furthermore, the percent freezing in DHL-SD animals was significantly lower than in DHL-NSD animals (p<0.001) with a 79.92% reduction in freezing on the 2^nd^ testing day. This indicates that prior exposure to CxFC did not facilitate the acquisition of contextual fear memory in the DH-lesioned animals subjected to SD. Additionally, SD prevented the formation of compensatory memory response in the DH-lesioned animals, resulting in impaired freezing expression even after providing the second conditioning. Overall these results emphasize the essential role of sleep in the induction of compensatory processes for contextual fear memory following DH loss.

### Effects of Pre-training NR2A Receptor Blockade in the Infralimbic Cortex (IL) on Compensatory Contextual Fear Memory After DH Loss

Here, we examined the role of NR2A receptors in the brain plasticity underlying the IL-dependent consolidation of contextual fear memory after DH loss. To investigate this, we blocked the activity of NR2A receptors by microinjecting PEAQX into DH-lesioned rats before their first CxFC training. Eight out of nine PEAQX microinjected animals with the IL region were included in the analysis (Fig. 4).

**Figure 4:**
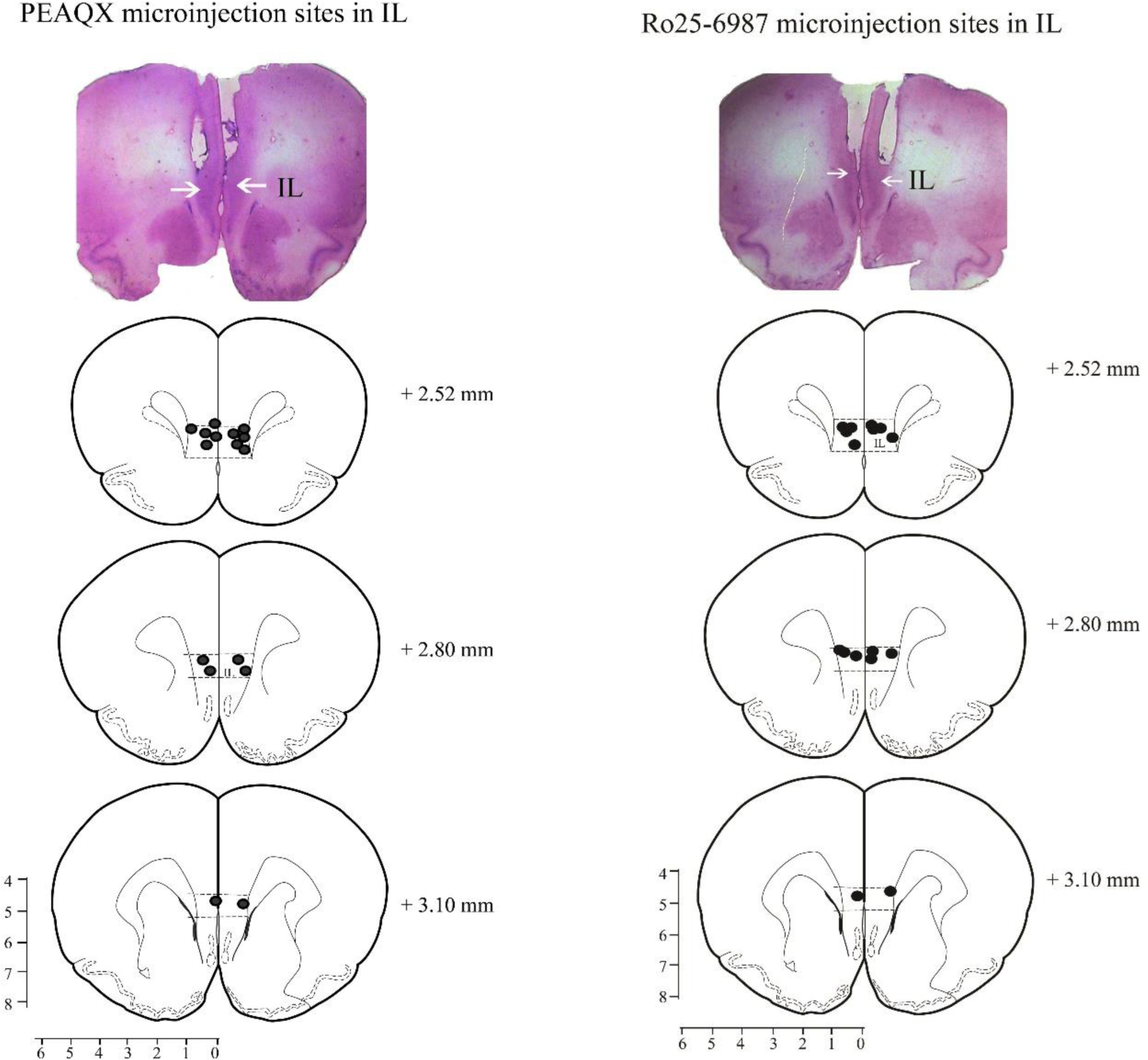
Cresyl violet-stained coronal section from rat brain with bilateral cannula tracks in IL and reconstruction diagram depicting the site of injection: NR2A-containing NMDA receptor antagonist (PEAQX) microinjection (n=8) and NR2B-containing NMDA receptor antagonist (Ro25-6981) microinjection (n=8). IL: Infralimbic cortex.

For the changes in percent freezing in the DH-lesioned animals with or without NR2A blockade, two-way ANOVA showed statistically significant interaction between drug treatment and day. The post hoc comparison demonstrated that DH-lesioned animals that received microinjections of either the NR2A-specific antagonist PEAQX or vehicle exhibited impaired freezing response on 1^st^ testing day (p<0.001; F _(8,114)_ = 11.60) (sample variance in DHL animals microinjected with vehicle in IL σ2 =143.21; DHL animals microinjected with PEAQX in IL σ2 = 223.39; Cohen’s d = 0.91, and power = 1.00 at the 0.05 α level) (Fig 5). After 2^nd^ training, DH-lesioned animals microinjected with vehicle displayed a robust increase in freezing response on 2^nd^ testing day, which was statistically significant (p<0.001, F _(8,114)_ = 11.60) compared to the 1^st^ testing day. Moreover, DH-lesioned animals that received microinjections of NR2A antagonist PEAQX also exhibited a significant increase (p<0.001) in freezing response on 2^nd^ testing day compared to 1^st^ testing day (Fig 5). The pre-training injection of the NR2A-specific antagonist did not impact the consolidation of compensatory contextual fear memory. These findings collectively indicate that NR2A-containing NMDA receptors are unlikely to play a substantial role in the IL-mediated consolidation of contextual fear in the absence of DH.

**Figure 5:**
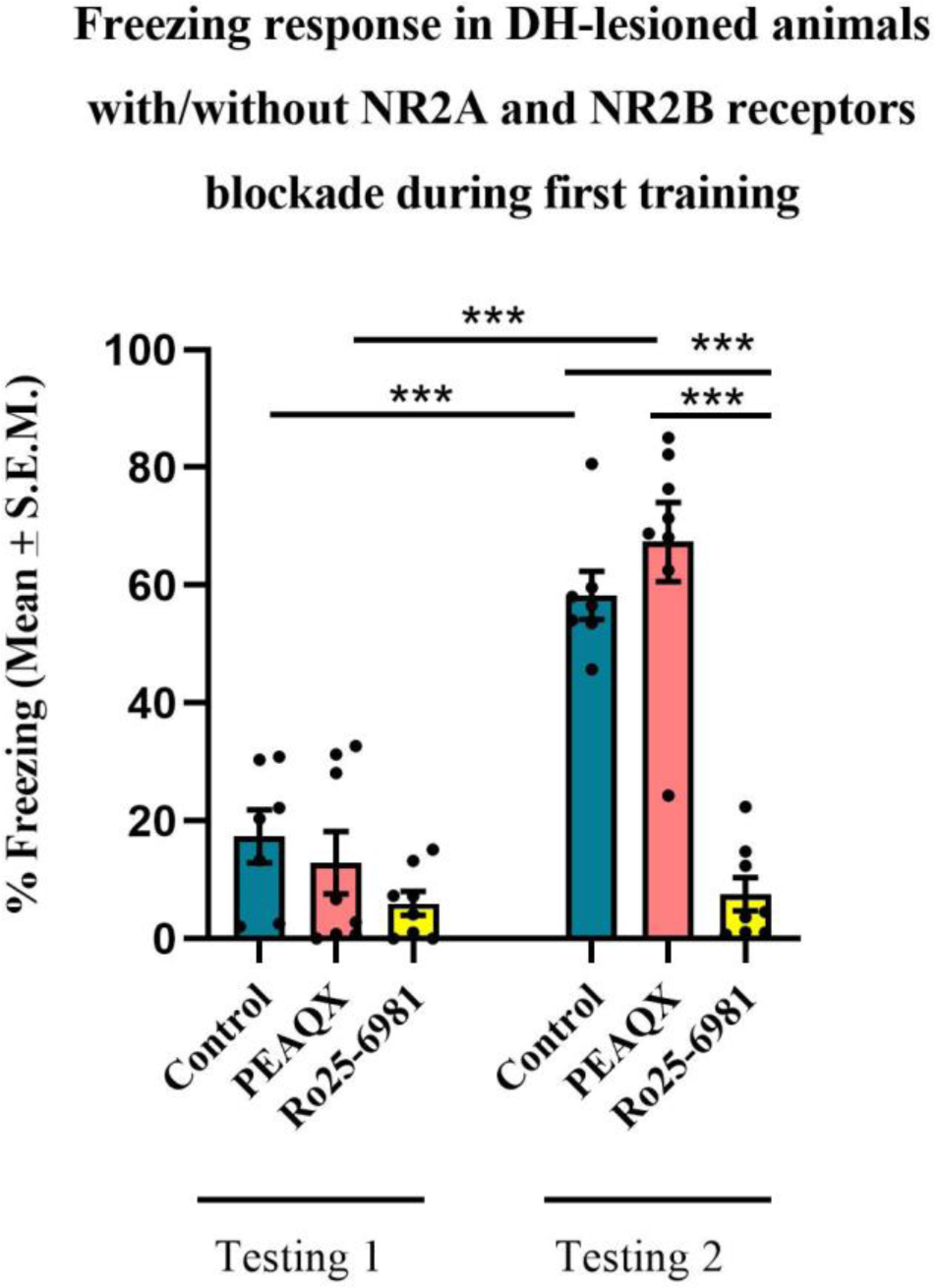
The role of NR2A and NR2B receptors in infralimbic cortex (IL)-mediated compensatory contextual fear memory following dorsal hippocampus (DH) loss: DH-lesioned rats received IL microinjections of the NR2A-specific antagonist PEAQX or vehicle before the first contextual fear conditioning (CxFC) training. Two-way ANOVA revealed a significant interaction between drug treatment and day. Both PEAQX and vehicle-treated DH-lesioned animals exhibited impaired freezing responses on the first testing day (p<0.001). Following the second training, both groups showed a significant increase in freezing response on the second testing day (p<0.001), indicating that pre-training NR2A blockade did not impact the consolidation of compensatory contextual fear memory. Additionally, DH-lesioned rats received IL microinjections of the NR2B-specific antagonist Ro25-6981 or vehicle before the first CxFC training. Both groups showed impaired freezing responses on the first testing day (p<0.001). However, following the second training, vehicle-injected DH-lesioned animals exhibited a significant increase in freezing response on the second testing day (p<0.001), whereas Ro25-6981-treated DH-lesioned animals showed significantly impaired freezing responses (p<0.001), indicating a critical role of NR2B subunits in the formation of compensatory contextual fear memory. Freezing response in DH-lesioned animals with/without NR2A and NR2B receptors blockade during first training

### Effects of Pre-training NR2B Receptor Blockade in the Infralimbic Cortex on Compensatory Contextual Fear Memory After DH Loss

To investigate the role of NR2B receptors in the neural plasticity underlying IL-dependent consolidation of contextual fear memory following DH loss, we utilized microinjections of Ro25-6981 to block NR2B receptor activity in DH-lesioned rats before the first CxFC training. Our analysis included 8 out of 9 animals that received successful Ro25-6981 microinjections within IL (Fig. 4).

Consistent with our prior findings, DH-lesioned animals subjected to microinjections of either the NR2A antagonist Ro25-6981 or vehicle exhibited impaired freezing responses on 1^st^ testing (Fig. 4). DH-lesioned animals injected with the vehicle displayed robust freezing responses on the 2^nd^ testing day, with a statistically significant increase (p<0.001, F _(8,114)_ = 11.60) compared to the 1^st^ testing day (Fig. 5). Conversely, DH-lesioned animals injected with the NR2B antagonist Ro25-6981 exhibited significant impaired freezing responses (p<0.001) on 2^nd^ testing day, with a significant reduction compared to their vehicle-injected counterparts (sample variance in DHL Ro25-6981 injected animals σ2 = 64.70; DHL vehicle-injected animals σ2 = 117.09; Cohen’s d = 4.68, and power = 1.00 at the 0.05 α level). Animals receiving the NR2B antagonist Ro25-6981 displayed an 87.02% reduction in freezing on 2^nd^ testing day compared to their vehicle-injected DH-lesioned counterparts, indicating the critical role of NR2B subunits in facilitating compensatory contextual fear memory formation in DH-lesioned animals.

Our overall results shed light on a distinctive role played by NMDA receptor subunits NR2A and NR2B in the consolidation of IL-dependent contextual fear memory in the absence of DH. The plasticity processes underlying IL-dependent consolidation of contextual fear memory after DH loss is selectively mediated by NR2B-containing NMDA receptors, while NR2A-containing receptors do not appear to be implicated in this particular process.

## Discussion

We observed that DHL-SD animals failed to compensate for DH loss even after multiple conditioning trials with a three-day gap. After the second conditioning, the DHL-SD group showed a significantly reduced freezing response on the 2^nd^ testing day (following 54 h of SD and 18 h of recovery) compared to the 1^st^ testing day. However, DHL-NSD animals displayed similar freezing responses on both testing days compared to the DHNL-NSD group. Additionally, our results suggest that NMDAR subunits NR2A and NR2B differentially regulate compensatory contextual fear formation without DH. Blocking NR2B subunits during the 1^st^ training inhibited compensatory fear memory formation, while blockade of NR2A subunits had no effect.

Previously, we showed that DH-lesioned rats develop a compensatory process for contextual fear conditioning when re-trained with a three-day gap. Reversible inactivation of the IL, during initial conditioning blocks this compensation. This suggests that the first contextual information reaching IL is vital for initiating the compensatory process, which attains functional maturity in about three days (Kant and Jha, 2019b, 2023). Our current study reinforces the importance of sleep and NR2B receptors, along with multiple conditioning trials and an active IL, for forming contextual fear memory without DH.

Many reports highlight the role of sleep in the development and reorganization of neuronal circuits. A study on cats revealed that new brain connections formed in the visual cortex of the animals during sleep developed twice the amount compared to the animals that remained awake (Frank *et al*., 2001; Jha *et al*., 2005b). Similarly, reorganization of the central olfactory system during sleep has been reported in mice. Here, they showed that newly generated adult-born neurons form connections with the pre-existing neural circuitry of the olfactory bulb. There occurs an incorporation/ elimination of newly formed neurons in the olfactory bulb during sleep and resting periods thereby reorganizing the circuit (Yamaguchi *et al*., 2013). Therefore, it may be possible that the development or reorganization of an alternative circuitry of contextual fear in IL is occurring predominantly during sleep. And, in the absence of sleep, the recruitment of this neural circuitry in IL might be getting impaired, thus preventing the overall compensatory process.

The role of mPFC sub-regions specifically IL and PL has been highlighted in the formation of contextual fear-conditioned memory in the absence of DH. Immediate early gene analysis showed that when contextual fear conditioning takes place during normal conditions (i.e. with an intact DH), the balance of activity between IL and PL is disrupted because of the decreased activity of IL. In situations, where the IL-PL micro-circuitry has to compensate for the loss of DH, a rearrangement in the balance of activity between IL and PL occurs (Zelikowsky *et al*., 2013). Furthermore, remodeling of cortical circuitry after learning at both the anatomical and functional levels has been reported during sleep, which is blocked by sleep deprivation (Dumoulin Bridi *et al*., 2015; Frank *et al*., 2001). Therefore, in our case, sleep deprivation may be leading to the disruption of balance between the activity of IL and PL and/or blockage of cortical circuitry remodeling that might be essential for compensating the loss of DH.

Sleep loss is known to negatively affect the functional connectivity of the prefrontal cortex with other brain areas. Additionally, recovery sleep proves to be beneficial for restoring the impaired functional connectivity of the prefrontal cortex region due to sleep loss (Durmer and Dinges, 2005). Sleep deprivation in humans poses a severe risk at the cognitive level thereby impairing the performance of simple day-to-day tasks like driving and machine-operating (Beattie *et al*., 2015). It has been reported that working memory, primarily mediated by the prefrontal cortex is particularly vulnerable to sleep restriction (Verweij *et al*., 2014). In our previous research, we demonstrated the crucial role of the IL in the formation of compensatory contextual fear memory. Specifically, when we inhibited IL in animals with damaged DH, we observed the complete disruption of the compensatory process (Kant and Jha, 2023). This finding implies that when contextual information reaches the IL for the first time, it initiates a reorganization of neural circuits, enabling the IL to compensate for the DH’s role in the expression of contextual fear responses. Interestingly, our investigations revealed that sleep deprivation, occurring during the temporal progression of these compensatory processes in the IL, hinders the compensation observed in DH-lesioned animals.

NMDARs play a significant role for the proper functioning of prefrontal region, such as facilitating LTP and working memory (Gilmartin *et al*., 2013; Zhao *et al*., 2005). Dysfunction of NMDARs within the prefrontal region leads to psychiatric disorders like schizophrenia (Marsman *et al*., 2011). A functional NMDAR is formed by an obligatory subunit NR1 and at least one NR2 subunit. According to the reports, NR2A and NR2B subunits are predominantly found within the prefrontal area (Loftis and Janowsky, 2003; Monyer *et al*., 1994). Additionally, NMDARs of different subunit combinations could be found in various brain regions during early development (Arrigoni and Greene, 2004; Monyer *et al*., 1994). However, compared with other cortical regions, mPFC is known to maintain a higher fraction of NR2B subunits during adulthood in rats (Wang *et al*., 2008). Our finding that the inactivation of NR2B and not NR2A subunits during the first contextual input inhibits the formation of compensatory fear memory, could be due to the prevalence of NR2B subunits over NR2A subunits in the IL region.

NMDAR subunits differentially regulate contextual fear memory across different brain regions. In the hippocampus, NR2A subunits are essential for contextual fear memory whereas NR2B subunits have a lesser impact (Gao *et al*., 2010b). Similarly, blocking NR2A unlike NR2B subunit in the prelimbic cortex severely impairs contextual fear response (Gilmartin *et al*., 2013). Conversely, contextual fear in the anterior cingulate cortex and amygdala, NR2B subunits predominantly mediate contextual fear (Rodrigues *et al*., 2001; Zhao *et al*., 2005). We have also observed that blocking NR2B subunits in the IL impairs the compensatory contextual fear response whereas blocking NR2A subunits do not have any impact. These findings emphasize the region-specific nature of NMDAR-mediated memory formation, closely associated with subunit composition. Furthermore, variable tyrosine phosphorylation states of NMDAR subunits between the hippocampus and cortex influence synaptic plasticity, giving rise to differences in long-term potentiation (LTP) and long-term depression (LTD) (Zhao *et al*., 2005). These changes in plasticity further emphasize the region-specific role of NMDARs in memory formation.

The functional differences of NMDARs are intricately related to their distinct subunit structures (Chen and Roche, 2007; Gilmartin *et al*., 2013). Additionally, NR2A and NR2B subunits differ in their channel properties, synaptic localization, and downstream signal transduction pathways (Chen and Roche, 2007; Zhao *et al*., 2005). NR2B, for instance, is predominantly found extrasynaptically, in contrast to NR2A subunits (Chen and Roche, 2007). Furthermore, the activation of NR2A subunits specifically triggers intracellular signaling pathways such as c-Fos and ERK1/2, whereas NR2B subunits primarily contribute to the membrane retention of pERK1/2 (Gao *et al*., 2010a). These distinctions suggest that the regulation of ERK signaling by NMDARs may be one of the molecular mechanisms contributing to the differential roles of their subunits in memory processes.

The hippocampus doesn’t work alone in encoding contextual fear memory; it works in concert with brain regions like the medial Prefrontal Cortex (mPFC), Retrosplenial Cortex (RSC), and Bed Nucleus of the Stria Terminalis (BST). When the primary DH network is compromised, subsidiary circuits gradually strengthen to consolidate fearful memories. Traumatic events associated with fear memories are pivotal in post-traumatic stress disorder (PTSD) development, causing patients to feel fear even in safe settings as they revisit memories. Normal fear memory processing is disrupted in PTSD, allowing fear memories to persist for years (Marchetta *et al*., 2023; van Marle). Our study suggests that this persistence could be due to fear memory encoding occurring in multiple brain regions. Understanding these compensatory pathways offers a new angle for developing PTSD therapies.

Our study contributes significantly to the understanding of fear memory processes, sleep, and NMDAR subunits. However, it is important to acknowledge certain limitations. Firstly, our experiments were conducted solely on male rats, despite the well-known sexual dimorphism in contextual fear conditioning. Including female rats would have provided a more comprehensive understanding of compensatory fear memory. Additionally, while we separately discussed the roles of sleep and NMDARs in fear memory processes, further research on their interplay, especially in compensatory fear memory, is needed. Exploring how sleep influences NMDAR-mediated memory consolidation could offer valuable insights into fear memory mechanisms.

In summary, our study highlights the significant impact of sleep on the alternative neural circuitry for contextual fear memory after DH loss. A 54-hour sleep deprivation period immediately following the first testing session was found to impair compensatory fear memory formation in DH-lesioned animals. The impact of sleep deprivation on compensatory processes is attributed to the loss of sleep itself, as observed in DHL-NSD animals compensating for DH loss through spaced training trials. Additionally, our findings highlight the importance of NR2B subunits in the IL for compensatory fear memory formation after DH loss. In contrast, NR2A subunits do not play a significant role in this process. Understanding the potential interplay between NMDARs and sleep in the absence of DH offers valuable insights into fear memory consolidation mechanisms and holds promise for novel therapeutic approaches targeting memory-related disorders.

## The Authors’ Contributions

DK has performed experiments, generated and analyzed data, and written the manuscript. SJ has conceived the idea, designed the experiments, analyzed the data, and finalized the manuscript.

