## Supplementary for "Sleep, NMDA Receptor Subunits, and the Compensatory Pathway: Understanding Contextual Fear Conditioning in the Absence of the Dorsal Hippocampus": Supplementary.docx

**Supplementary information**

Partial Sleep Deprivation and Compensatory Memory Formation: We aimed to investigate sleep's role in the formation of compensatory neural circuitry for contextual fear memory in the absence of the DH. Previous research showed that DH-lesioned animals, when re-trained three days after initial conditioning, recover contextual fear responses similar to non-lesioned animals. This three-day (72-hour) interval between multiple conditioning is critical for the alternative circuitry to compensate for DH's role in memory consolidation.

To explore the direct influence of sleep during this period, we conducted sleep deprivation (SD) experiments. SD is known to impair both memory acquisition and consolidation processes. We tested three different SD protocols (supplementary fig. 1):

1. After baseline recording, animals were subjected to 60 h SD + 12 h recovery sleep followed by CxFC (60SD12R-C): which significantly (p<0.001, F_(8,44)_ = 17.63) impaired contextual fear memory acquisition as suggested by decreased percent freezing on the testing day compared to non-sleep-deprived (NSD) animals (sample variance in NSD animals σ2 = 22.36; 60SD12R-C animals σ2 = 8.18; Cohen’s d = 2.46, and power = 1.00 at the 0.05 α level) (supplementary fig. 2).
2. After baseline recording, animals were subjected to 54 h SD + 18 h recovery sleep followed by CxFC (54SD18R-C): which also resulted in significant memory impairment as suggested by decreased percent freezing (p<0.001, F_(8,44)_ = 17.63) on the testing day compared to NSD animals (sample variance in NSD animals σ2 = 22.36; 54SD18R-C animals σ2 = 9.17; Cohen’s d = 2.86, and power = 1.00 at the 0.05 α level) (supplementary fig. 2).
3. After baseline recording, animals were subjected to first CxFC, followed by 54 h SD + 18 h recovery sleep, and second CxFC (C-54SD18R-C): Animals exhibited robust freezing response as suggested by significantly increased percent freezing (p<0.001, F_(8,44)_ = 17.63) on second testing compared to baseline day (sample variance in C-NSD-C animals σ2 = 22.31; C-54SD18R-C animals σ2 = 24.51; Cohen’s d = 0.23, and power = 1.00 at the 0.05 α level). This indicated that prior CxFC exposure facilitated memory acquisition in sleep-deprived animals (supplementary fig. 2).

Since the third protocol (C-54SD18R-C) facilitated effective learning under SD conditions and aligned with our established protocol (two CxFC sessions separated by 3 d), we selected this protocol for experiment I.

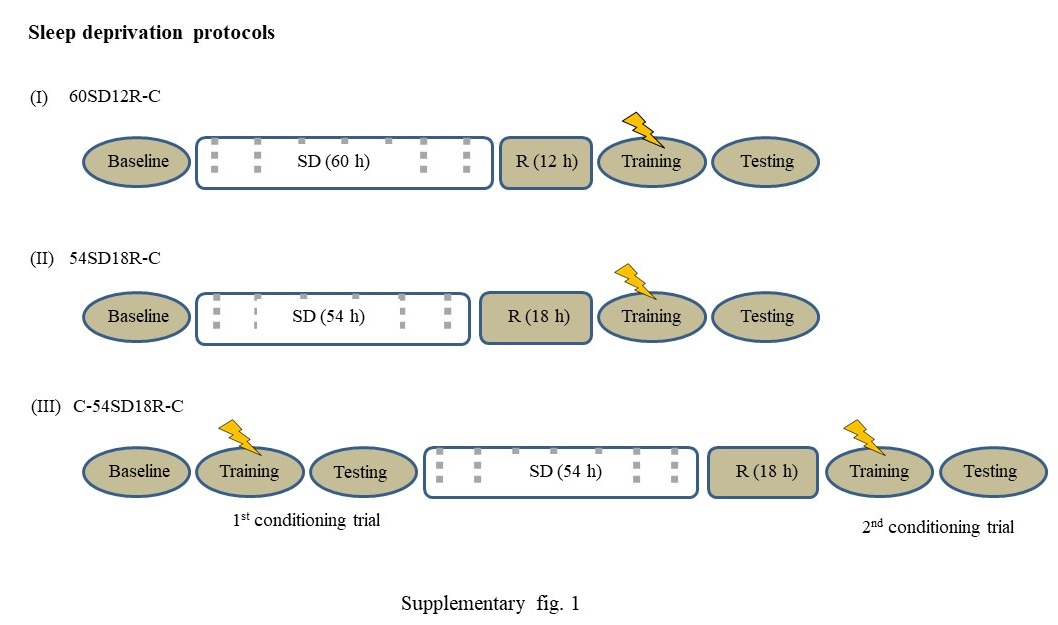

**Supplementary Figure 1:** Standardization of Sleep Deprivation Protocol.

Protocol I: Animals were subjected to 60 hours of sleep deprivation followed by 12 hours of recovery sleep, and then underwent contextual fear conditioning (CxFC). This protocol is denoted as 60SD12R-C.

Protocol II: Animals experienced 54 hours of sleep deprivation followed by 18 hours of recovery sleep before undergoing CxFC. This protocol is referred to as 54SD18R-C.

Protocol III: Animals first underwent CxFC, followed by 54 hours of sleep deprivation and 18 hours of recovery sleep, and then underwent a second CxFC. This protocol is designated as C-54SD18R-C.

Here, SD: sleep deprivation, R: recovery, C: contextual fear conditioning.

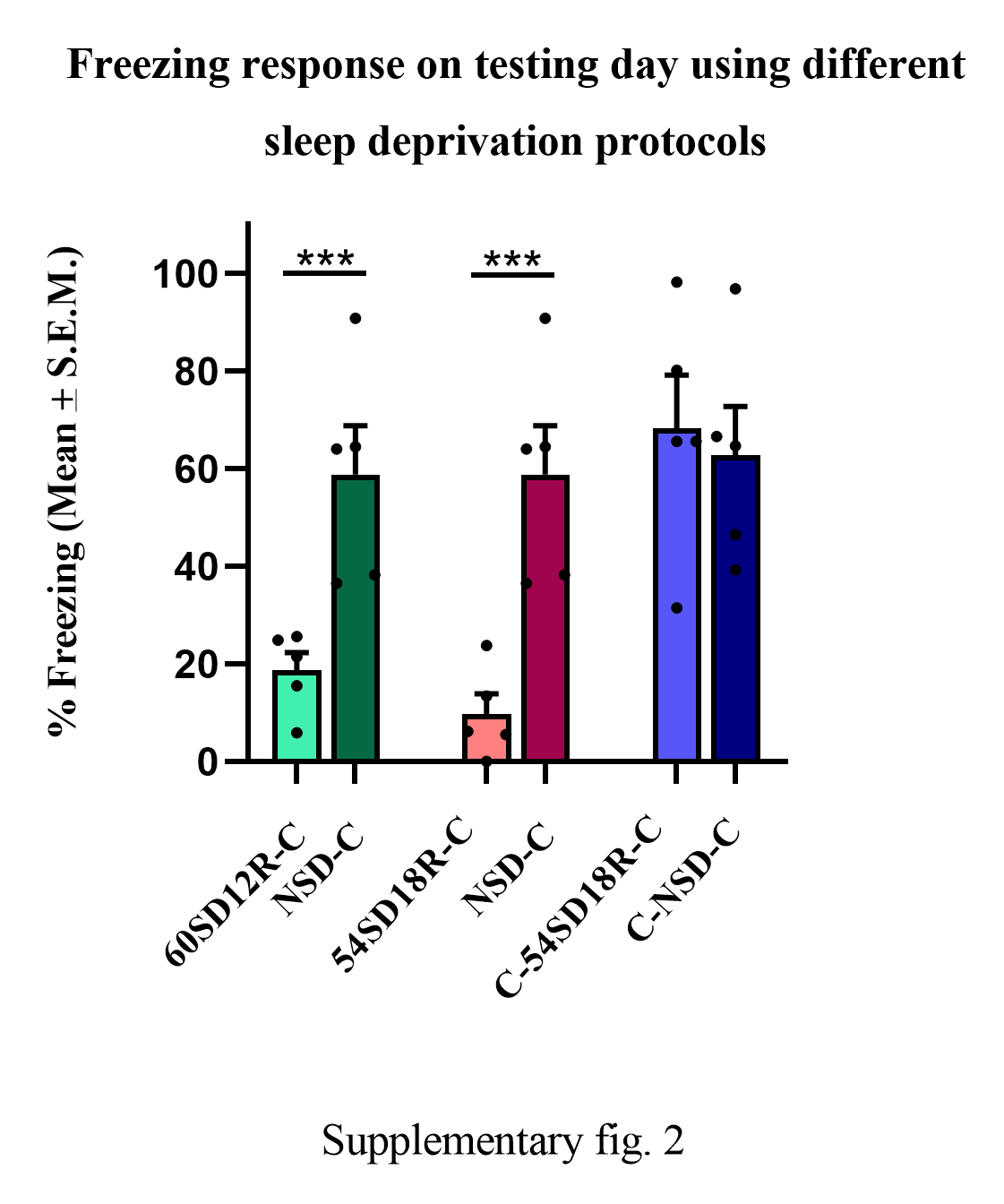

**Supplementary Figure 2:** Effects of Sleep Deprivation and Recovery Sleep on Contextual Fear Memory Acquisition

Protocol I (60SD12R-C): Animals (n=5) were subjected to 60 hours of sleep deprivation followed by 12 hours of recovery sleep, then contextual fear conditioning (CxFC). This protocol significantly impaired contextual fear memory acquisition, as inferred by a decreased percentage of freezing on the testing day compared to non-sleep-deprived (NSD) animals (p < 0.001.

Protocol II (54SD18R-C): Animals (n=5) were subjected to 54 hours of sleep deprivation followed by 18 hours of recovery sleep, then CxFC. This also resulted in significant memory impairment, indicated by a decreased freezing percentage on the testing day compared to NSD animals (p < 0.001).

Protocol III (C-54SD18R-C): Animals (n=5) first underwent CxFC, followed by 54 hours of sleep deprivation and 18 hours of recovery sleep, and then a second CxFC. These animals exhibited a robust freezing response, with a significantly increased freezing percentage on the second testing day compared to the baseline day (p < 0.001). This indicated that prior CxFC exposure facilitated memory acquisition in sleep-deprived animals.

*** denotes Tukey *p* < 0.001. Error bars represent the standard error mean (S.E.M.).
